# Hierarchical Uncertainty Transformation During Multimodal Comprehension

**DOI:** 10.64898/2026.10.29.762725

**Authors:** Wanying Zhao, Jiawang Yang, Zhouyi Li, Yi Du

## Abstract

Human communication requires combining information from different modalities even when each source is incomplete or ambiguous. Yet how uncertainty is coordinated as distinct semantic representations interact and converge remains unclear. Here we propose a hierarchical uncertainty transformation framework and test it across four gesture–speech experiments combining behavioral measures, information-theoretic modelling, and electroencephalography. Rather than being uniformly reduced as information accumulates, uncertainty changed its computational form across three interdependent levels: uncertainty within modality-specific semantic representations, bidirectional but asymmetric changes during cross-modal interaction, and uncertainty associated with multisensory convergence. These computational distinctions were accompanied by differentiated but temporally overlapping neural associations. Modality-specific uncertainty was expressed relatively early, whereas cross-modal transformation and convergence were most prominently associated with activity in the N400 range. Critically, both the neural expression and statistical coupling of these computational quantities varied with the semantic information available from gesture and speech. Together, these findings show that multimodal comprehension involves a context-dependent reorganization of uncertainty across interacting representations, providing a computational framework for understanding how coherent meaning emerges from heterogeneous and uncertain information.

## Introduction

Human communication depends on the ability to construct coherent meaning from information distributed across modalities. In face-to-face interaction, speech and co-speech gestures jointly convey meaning despite differing in representational format, temporal dynamics and informational structure^1^. Comprehending such input requires heterogeneous and temporally asynchronous signals to be integrated in real time^2^, even though information arriving through either modality may be incomplete or ambiguous^3,4^. Multimodal comprehension must therefore solve a fundamental computational problem: how can information distributed across distinct representational streams be transformed into a coherent interpretation?

A key feature of semantic processing is that incomplete or ambiguous signals can support multiple competing interpretations^5^. Semantic representations can therefore be conceptualized as probability distributions over possible interpretations^6,7^ rather than as discrete meanings. Neural responses during language comprehension exhibit graded activation^8,9^ and competition among candidate meanings^10,11^, while behavioral responses are sensitive to probabilistic structure arising from frequency-dependent recognition, contextual disambiguation, and expectation-based processing^12^. The uncertainty associated with a semantic representation can thus be understood in terms of how probability is distributed across its possible interpretations. Multimodal communication must consequently operate not only on the semantic content, but also on the uncertainty associated with that content^13^.

Computational accounts provide powerful perspectives on how information from multiple sensory sources is integrated. Prominent approaches include Bayesian causal inference and predictive processing. Bayesian causal inference models formalize how uncertain sensory signals are integrated while accounting for their inferred causal structure^14,15^, whereas predictive-processing accounts characterize perception in terms of predictions and the updating of internal representations through prediction error^16–20^. These approaches have accounted for a broad range of behavioral and neural phenomena but have primarily focused on the principles governing evidence combination or representational updating. Less clear is how uncertainty itself changes across different levels of multimodal processing. This question is particularly important because multisensory signals are processed in hierarchically organized systems^21,22^, unfold over distinct temporal scales^23^, and interact through multiple processing stages^24,25^. Under these conditions, a reduction in uncertainty at one level need not entail an equivalent reduction at another, raising the possibility that multimodal integration involves changes in the form, rather than simply the amount, of uncertainty.

Here we propose a hierarchical uncertainty transformation framework for multimodal semantic integration. Information theory provides a unified quantitative framework for characterizing uncertainty and changes in uncertainty across representations^26,27^. We distinguish three computational levels: modality-specific semantic uncertainty, reflecting uncertainty over possible interpretations within each modality; cross-modal uncertainty transformation, reflecting changes in modality-specific uncertainty in the presence of information from the other modality; and multisensory uncertainty convergence, reflecting the organization of uncertainty as modality-specific information converges within an integrated semantic representation (**Fig. 1c**). We hypothesize that these levels capture distinct but interdependent computational forms of uncertainty rather than fixed temporal stages of integration. Accordingly, their neural expressions may be temporally differentiated yet overlapping and should vary with the semantic information available from each modality.

**Fig. 1.**
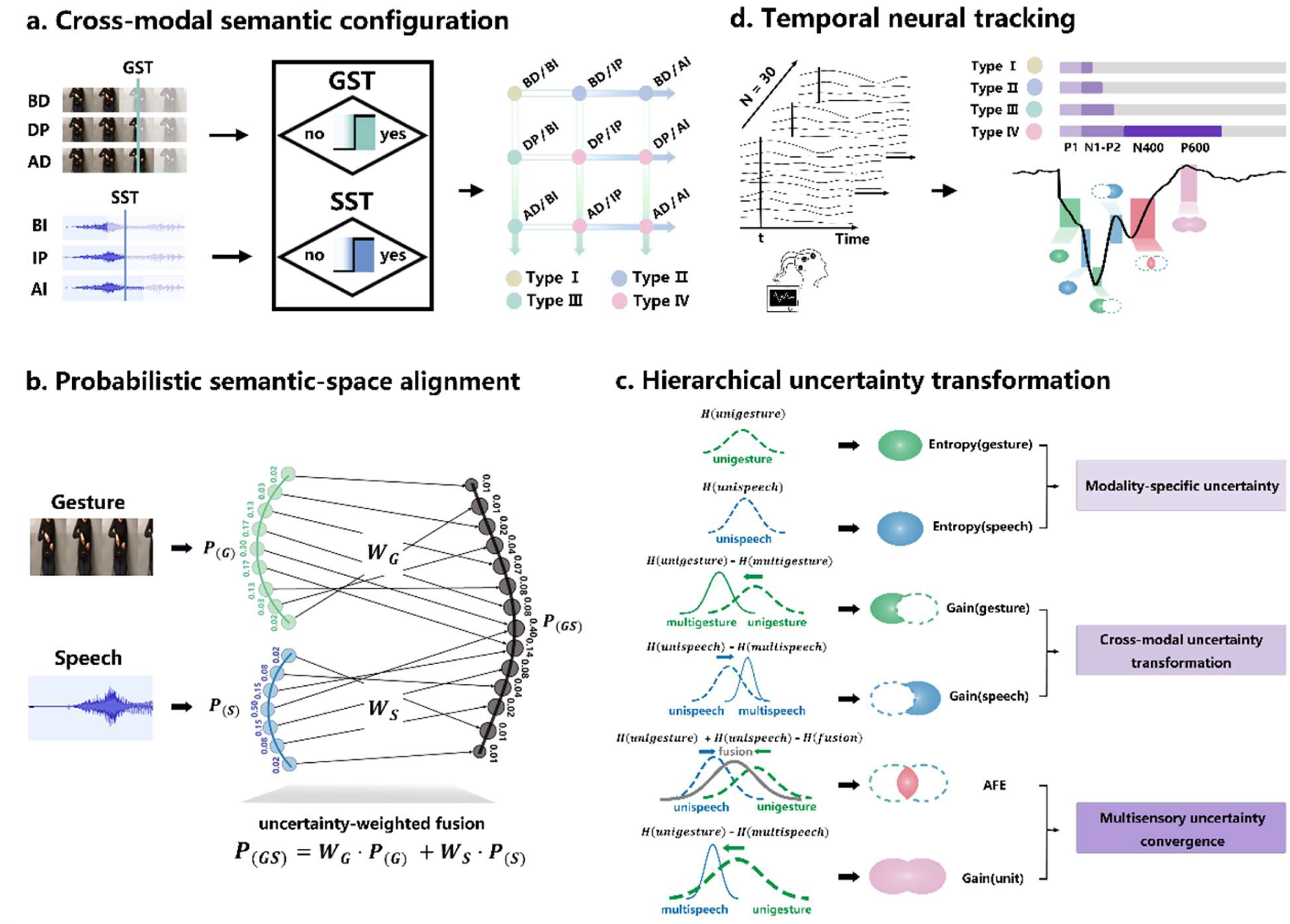
Experimental and computational framework for hierarchical uncertainty transformation during multimodal semantic integration. **a,** Cross-modal semantic configuration. For each modality, gesture and speech fragments were defined relative to independently determined semantic-identification thresholds and sampled at pre-threshold, threshold and post-threshold semantic-specification states. Crossing the three gesture and three speech states yielded nine fragment conditions spanning different combinations of modality-specific semantic information. For analyses of broader cross-modal integration contexts, these nine conditions were further grouped into four cross-modal semantic configurations according to whether each modality had reached its semantic-identification threshold: Type I, both modalities pre-threshold; Type II, gesture pre-threshold and speech threshold-reached; Type III, gesture threshold-reached and speech pre-threshold; and Type IV, both modalities threshold-reached. b, Probabilistic semantic-space alignment. Naming responses to gesture and speech fragments were converted into modality-specific semantic probability distributions. The modality-specific semantic vocabularies were mapped onto a shared vocabulary in which identical semantic labels occupied the same dimension, while modality-specific labels were retained and assigned zero probability in the other modality. This dimension-wise alignment yielded comparable gesture and speech probability distributions within a shared probabilistic semantic space, which were subsequently combined using uncertainty-weighted fusion to obtain a fused semantic probability distribution. c, Hierarchical uncertainty transformation. Three computational levels were used to characterize distinct aspects of uncertainty transformation. Level 1 quantified modality-specific semantic uncertainty using gesture and speech entropy. Level 2 quantified cross-modal changes in modality-specific uncertainty using gesture gain and speech gain, defined as the change in entropy from the unisensory to the corresponding multisensory context. Level 3 quantified multisensory uncertainty convergence using aligned fusion entropy (AFE) and unit gain. These levels represent distinct computational quantities rather than discrete serial processing stages. d, Temporal neural tracking. EEG was used to characterize the temporal neural expression of the three computational levels and to determine how their neural associations varied across cross-modal semantic configurations. Time-resolved representational and computational-EEG analyses were used to identify neural signatures associated with modality-specific uncertainty, cross-modal uncertainty transformation and multisensory uncertainty convergence. BD, before gesture discrimination point; DP, gesture discrimination point; AD, after gesture discrimination point; BI, before speech identification point; IP, speech identification point; AI, after speech identification point; GST, gesture semantic threshold; SST, speech semantic threshold; AFE, aligned fusion entropy; EEG, electroencephalography.

We tested this framework in gesture-speech communication by combining behavioral measures of semantic interpretations, information-theoretic modelling and time-resolved electroencephalography (EEG). We independently varied gesture and speech fragments relative to empirically determined semantic-identification thresholds, thereby manipulating the semantic information available from each modality and generating four cross-modal semantic configurations in which neither, either or both modalities had reached threshold (**Fig. 1a**). Gesture and speech naming distributions were aligned within a shared probabilistic semantic space and combined using uncertainty-dependent modality weights, enabling uncertainty and its transformation to be quantified within a common representational framework (**Fig. 1b**). The three levels showed distinct computational profiles and differentiated but overlapping neural associations whose temporal expression varied across cross-modal semantic configurations (**Fig. 1d**). Cross-level analyses further revealed statistical dependencies between cross-modal uncertainty transformation and multisensory convergence, with these dependencies also varying across configurations. Together, these findings support a hierarchical uncertainty transformation account in which multimodal semantic integration changes the computational form of uncertainty across distinguishable but interdependent levels rather than simply reducing uncertainty as information accumulates.

## Results

We investigated whether multimodal semantic integration can be characterized as a hierarchical transformation of uncertainty rather than a uniform reduction of uncertainty across modalities. We distinguished three computational levels: modality-specific semantic uncertainty, cross-modal uncertainty transformation and multisensory uncertainty convergence. Semantic uncertainty was quantified from the probability distributions of naming responses using Shannon entropy^28^. Cross-modal transformation was quantified as changes in modality-specific entropy between unisensory and multisensory contexts^29^, whereas multisensory convergence was characterized using two complementary measures derived from the aligned gesture and speech probability distributions: aligned fusion entropy (AFE) and unit gain. We first established the computational profiles of these measures, then examined whether they show differentiated neural associations, whether their neural expression varied with the joint semantic specification of gesture and speech, and whether the computational levels were statistically interconnected.

### Hierarchical transformation from modality-specific uncertainty to multisensory convergence

#### *Level 1:* Modality-specific uncertainty varies with semantic specification

At the first computational level, we examined how modality-specific semantic uncertainty varied across the three semantic specification states defined relative to the independently determined semantic-identification threshold: pre-threshold, threshold and post-threshold. Within the hierarchical uncertainty-transformation framework, this level characterizes the modality-specific uncertainty available for subsequent cross-modal transformation. We therefore expected uncertainty to vary systematically with increasing semantic specification in each modality. Entropy was calculated separately for gesture and speech from the empirical probability distributions of naming responses, with higher values indicating greater dispersion across possible semantic interpretations.

Semantic entropy decreased with increasing semantic specification in both modalities. Gesture entropy declined significantly across semantic-specification states (*F*(2, 54) = 9.577, *p* < 0.001, η^2^ = 0.262, **Fig. S1a1**), as did speech entropy (*F*(2, 54) = 35.481, *p* < 0.001, η^2^ = 0.568, **Fig. S1a2**). In both modalities, entropy was highest in the pre-threshold state and lower once the semantic-identification threshold was reached. These results establish the modality-specific uncertainty structure on which subsequent cross-modal transformation operates and confirm that the fragment manipulation systematically varies semantic uncertainty while preserving continuous stimulus-level variation within each semantic-specification state.

#### *Level 2:* Cross-modal uncertainty transformation is bidirectional and asymmetric

At the second computational level, we examined how modality-specific uncertainty changed when information from the other modality became available. If multimodal integration involves cross-modal transformation rather than a uniform reduction of uncertainty, these changes should depend on the semantic specification of the two modalities and need not be equivalent for gesture and speech. Gesture gain and speech gain quantified changes in modality-specific entropy from the unisensory to the corresponding multisensory context, with positive values indicating reduced uncertainty.

Gesture gain varied as a function of both gesture and speech semantic specification states (gesture: *F*(1.49, 26.81) = 20.168, η^2^ = 0.528; speech: *F*(1.41, 25.28) = 38.429, η^2^ = 0.681; both *p* < 0.001, **Fig. S1b**). Larger gesture gains were observed when gesture remained relatively uncertain and speech was more semantically specified. Speech gain likewise varied with gesture semantic specification (*F*(1.85, 33.22) = 17.279, *p* < 0.001, η^2^ = 0.490, **Fig. S1c1**) but showed no reliable modulation by speech semantic specification state (*p* = 0.119, **Fig. S1c2**).

Thus, cross-modal information altered uncertainty in both modality-specific representations, but the transformation profiles differed between gesture and speech. The pattern was consistent with relatively more semantically specified information in one modality preferentially constraining uncertainty in the less specified modality, although this influence was asymmetric across gesture and speech. These bidirectional but asymmetric changes are inconsistent with a uniform reduction of modality-specific uncertainty and instead support the view that cross-modal information transforms modality-specific uncertainty in a structured manner.

#### *Level 3:* Multisensory uncertainty convergence comprises complementary computational measures

At the third computational level, we examined how cross-modal transformation culminated in a form of multisensory convergence that could not be reduced to modality-specific entropy changes alone. If hierarchical uncertainty transformation extends beyond changes within gesture and speech representations, measures defined over the integrated semantic system should show computational profiles that differ from modality-specific gains. We therefore characterized convergence using two complementary measures: AFE, which quantified the change in uncertainty associated with uncertainty-weighted fusion, and unit gain, which quantified the change in uncertainty from the initial gesture representation to the subsequent speech representation in the multisensory context.

AFE varied significantly with the semantic specification states in both modalities (speech: *F*(1.61, 28.96) = 35.702, η^2^_p_ = 0.665; gesture: *F*(1.45, 26.07) = 49.118, η^2^ = 0.732; both *p* < 0.001, **Fig. S1d**). AFE was highest under pre-threshold conditions and decreased as semantic specification increased. Unit gain showed a different profile, varying significantly with speech semantic specification (*F*(1.84, 33.09) = 17.426, *p* < 0.001, η^2^ = 0.492, **Fig. S1e2**), and increasing after speech reached its semantic-identification threshold.

The divergent profiles of AFE and unit gain indicate that multisensory convergence cannot be described as a simple continuation of modality-specific uncertainty reduction. Instead, the two measures capture complementary aspects of uncertainty at the integrated level. Together with the bidirectional and asymmetric gain effects, these findings support a hierarchical transformation account in which uncertainty changes form across computational levels—from modality-specific uncertainty, through cross-modal transformation, to multisensory convergence.

### Neural dynamics differentiate computational levels of uncertainty transformation

Having established the three computational levels, we next asked whether these computational components were differentiated in their neural expression. If they capture distinguishable components of a hierarchical transformation process rather than alternative descriptions of a single uncertainty-reduction signal, their corresponding information-theoretic measures should exhibit differentiated neural associations. We therefore first used time-resolved full-factorial representational similarity analysis (RSA) to identify neural representations associated with gesture semantic specification, speech semantic specification and semantic congruency, and then related activity within the corresponding RSA-defined clusters to the continuous information-theoretic measures.

Gesture semantic specification was represented at multiple latencies, including early P1 activity (0-100 ms: *β* = 3.36, *p* < 0.001), the N1-P2 complex (150-250 ms: *β* = 1.82, *p* < 0.001), and later N400-range intervals (336-406 ms, *β* = 0.77; 446-500 ms, *β* = 0.71; both *p* < 0.001, **Fig. 2a green line**). Within the early P1 gesture-related RSA cluster, ERP amplitude was negatively correlated with gesture entropy over frontal-central channels (*r* = -0.19 to -0.32, false discovery rate (FDR)-corrected *p* < 0.017, **Fig. 2b Table Cell_1,1_**).

**Fig. 2.**
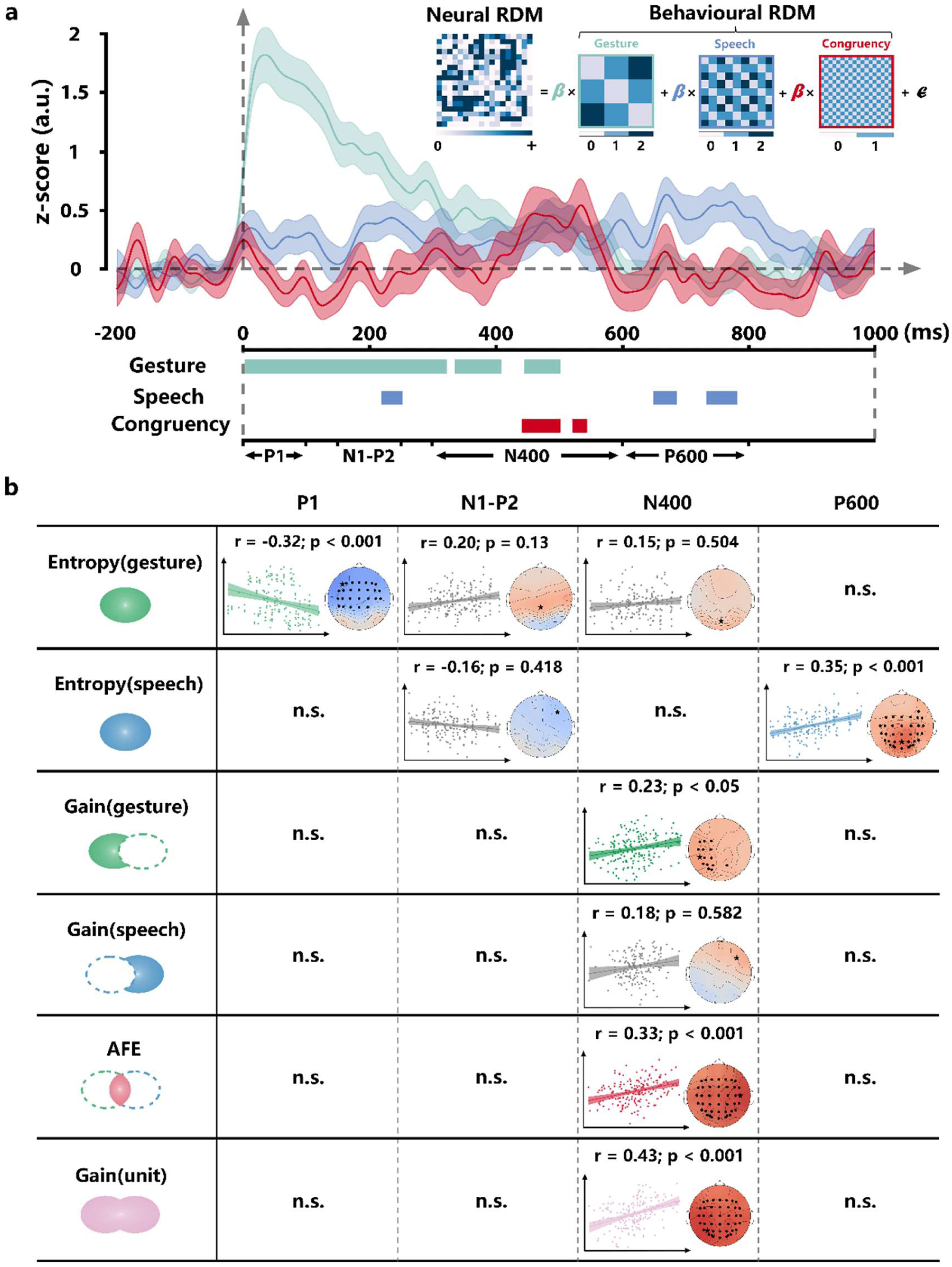
Temporally differentiated neural signatures across the uncertainty-transformation hierarchy. **a,** Time-resolved full-factorial representational similarity analysis (RSA) showing baseline-corrected, standardized regression coefficients for gesture semantic specification, speech semantic specification and semantic congruency. At each time point, an 18 × 18 neural representational dissimilarity matrix (RDM) was constructed across experimental conditions and simultaneously regressed against model RDMs representing the three experimental dimensions; regression coefficients therefore quantify the unique correspondence between each model RDM and the neural representational geometry. Gesture semantic specification was represented during the P1 (0-100 ms), N1-P2 (150-250 ms) and N400 ranges (336-406 and 446-500 ms); speech semantic specification during the N1-P2 (220-250 ms) and P600 ranges (650-780 ms); and semantic congruency prominently during the N400 range (442-542 ms). Shaded areas indicate ±1 s.e.m. Insets show representative model and neural RDMs. **b,** Associations between continuous information-theoretic measures and ERP activity within the corresponding RSA-defined clusters. Gesture entropy was negatively correlated with activity over frontal-central electrodes within the gesture-related cluster, whereas speech entropy was positively correlated with posterior activity within the speech-related cluster. Within the semantic-congruency-related cluster, the congruency difference wave was associated with gesture gain, AFE and unit gain, but not reliably with speech gain or modality-specific entropy. Correlations were calculated using Spearman’s rank correlation and corrected for multiple comparisons using the false-discovery rate (FDR). AFE, aligned fusion entropy; ERP, event-related potential; RDM, representational dissimilarity matrix; RSA, representational similarity analysis.

Speech semantic specification showed a different temporal profile, with significant RSA effects during the N1-P2 interval (220-250 ms, *β* = 0.43, *p* < 0.001) and late P600 stages (650-780 ms, *β* = 0.58 -0.66, *p* < 0.001, **Fig. 2a blue line**). Speech entropy was associated with ERP amplitude over posterior electrodes during the P600 window (*r* = 0.17 -0.35, *p* < 0.042, **Fig. 2b Table Cell_2,4_**). Thus, modality-specific semantic uncertainty was associated with modality-related neural responses, but its temporal expression differed between gesture and speech.

Neural sensitivity to semantic congruency emerged prominently in the N400 range (442-542 ms, *β* = 0.55 - 0.56, *p* < 0.001, **Fig. 2a red line**), identifying a temporal interval associated with gesture-speech semantic interaction. Within this semantic-congruency-related RSA cluster, the congruency ERP difference was correlated with gesture gain over left central-posterior channels (*r* = 0.18 - 0.23, *p* < 0.048, **Fig. 2b Table Cell_3,3_**), linking N400 activity to cross-modal uncertainty transformation. No reliable association with speech gain was observed in this analysis (*r* = 0.18, *p* = 0.582, **Fig. 2b Table Cell_4,3_**).

The same N400-range activity (442-542 ms) was also correlated with the multisensory convergence measures. Across all scalp channels, significant correlations were observed for AFE (*r* = 0.24 - 0.33, *p* < 0.002, **Fig. 2b Table Cell_5,3_**) and unit gain (*r* = 0.16 - 0.43, *p* < 0.048, **Fig. 2b Table Cell_6,3_**). No reliable association with modality-specific entropy was detected within the semantic-congruency-related cluster. Complementary analyses using predefined ERP time windows and regions of interest (ROIs) showed a comparable pattern of associations across modality-specific uncertainty, cross-modal uncertainty transformation and multisensory convergence (**Fig. S2** and **Supplementary Results**).

Together, these results provide neural differentiation across the computational hierarchy. Modality-specific semantic uncertainty was preferentially associated with modality-related neural responses at distinct temporal intervals, whereas cross-modal uncertainty transformation and multisensory uncertainty convergence were more closely associated with neural activity related to semantic congruency, particularly in the N400 range. Importantly, however, these associations were temporally differentiated but partially overlapping. The computational hierarchy therefore did not map onto a fixed sequence of discrete neural stages; rather, distinct forms of uncertainty transformation showed distinguishable but partially overlapping neural expression.

### Cross-modal semantic configuration shapes the neural expression of uncertainty transformation

Having established neural differentiation across the computational hierarchy, we next tested whether its expression was invariant or depended on the semantic information available from the two modalities. If hierarchical uncertainty transformation is organized by cross-modal semantic context, the neural expression of its computational components should vary with the joint semantic specification of gesture and speech rather than follow a fixed temporal sequence.

The nine fragment conditions were therefore grouped into four broader cross-modal semantic configurations according to whether each modality had reached its independently determined semantic-identification threshold: Type I, gesture pre-threshold & speech pre-threshold; Type II, gesture pre-threshold & speech threshold-reached; Type III, gesture threshold-reached & speech pre-threshold; and Type IV, gesture threshold-reached & speech threshold-reached (**Fig. 1a**). Continuous information-theoretic measures were retained within each configuration.

### Configuration-dependent neural expression of modality-specific semantic uncertainty

We first examined whether the neural expression of Level 1 uncertainty depended on the semantic configuration in which it occurred. Gesture entropy showed early P1-N1-P2 associations across all four configurations (all *p* < 0.001; **Fig. 3a**), indicating that gesture uncertainty was neurally expressed even when little semantic information was available from either modality. Later associations, however, varied with cross-modal semantic configuration. N400-range effects emerged selectively when speech was threshold-reached (Types II and IV, *p* < 0.001), and extended P2-N400 clusters were observed when gesture itself was threshold-reached (Types III and IV, *p* < 0.026).

**Fig. 3.**
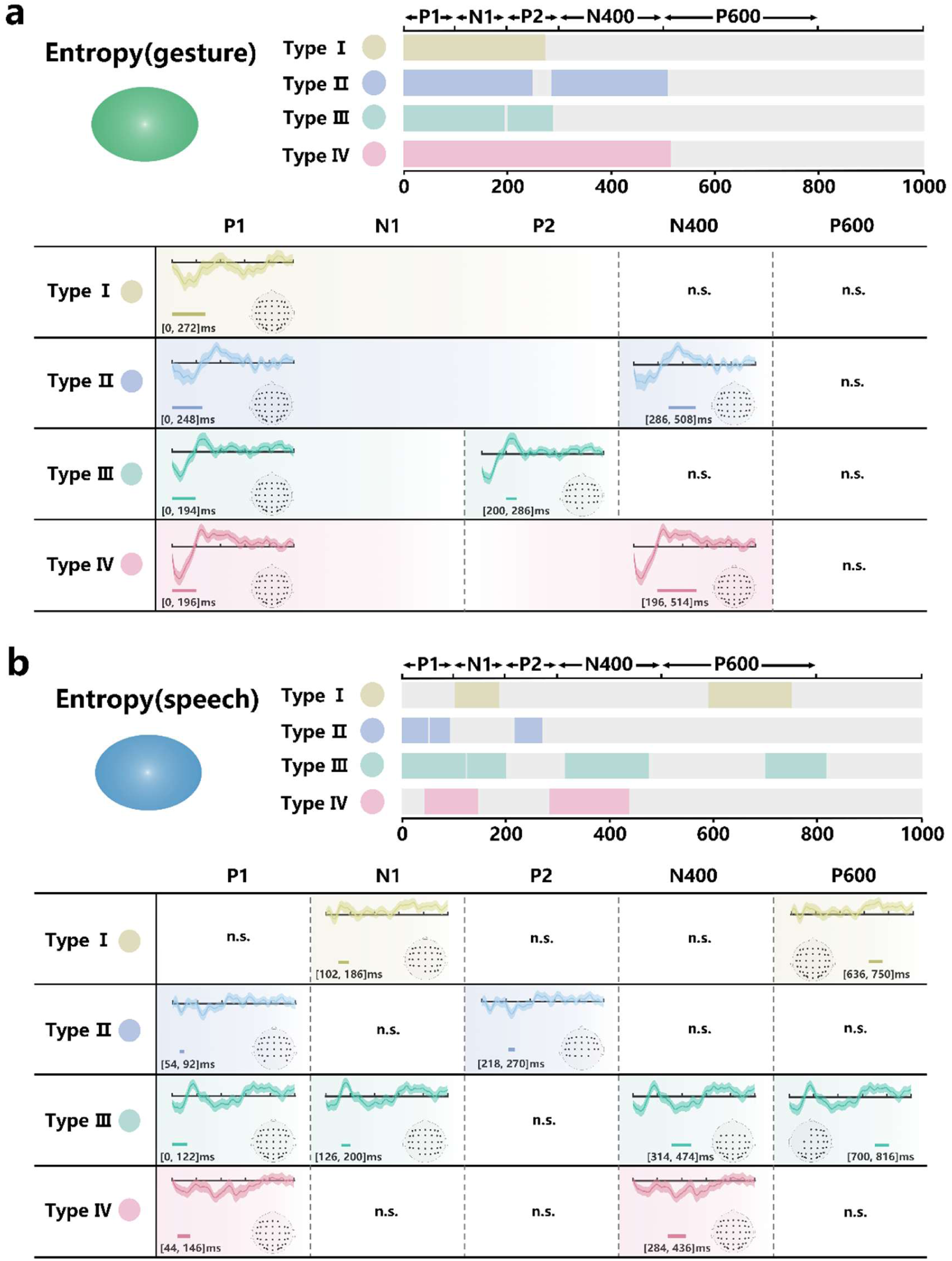
Cross-modal semantic configuration shapes the neural expression of modality-specific semantic uncertainty. **a,b**, Spatiotemporal associations between ERP activity and continuous gesture entropy (**a**) or speech entropy (**b**) were examined separately within four cross-modal semantic configurations. Type I comprised conditions in which both gesture and speech were pre-threshold; Type II, gesture pre-threshold and speech threshold-reached; Type III, gesture threshold-reached and speech pre-threshold; and Type IV, both modalities threshold-reached. Entropy was calculated from the probability distribution of naming responses for each modality, with higher values indicating greater semantic uncertainty. Participant-level Spearman correlation coefficients were tested against zero using two-tailed spatiotemporal cluster-based permutation tests with 1,000 permutations. **a**, Gesture entropy showed a temporally extended pattern, with P1-N1-P2 associations across all four configurations and later associations that varied with cross-modal semantic configuration. N400-range effects were observed when speech was threshold-reached (Types II and IV), whereas extended P2-N400 effects occurred when gesture was threshold-reached (Types III and IV). **b**, Speech entropy showed a strong configuration-dependent temporal profile. Early associations shifted from the N1 range when both modalities were pre-threshold (Type I) to the earlier P1 range when either modality was threshold-reached (Types II-IV). At later latencies, N400 associations occurred when gesture was threshold-reached (Types III and IV), whereas P600 associations occurred when speech remained pre-threshold (Types I and III). ERP, event-related potential.

Speech entropy showed a more strongly configuration-dependent temporal profile **(Fig. 3b)**. Early associations shifted from the N1 range when both modalities were pre-threshold (Type I, *p* = 0.031) to the earlier P1 range when either modality had reached its semantic-identification threshold (Types II-IV, *p* ≤ 0.017). At later latencies, N400 associations emerged when gesture was threshold-reached (Types III and IV, *p* < 0.001), whereas P600 associations were observed when speech remained pre-threshold (Types I and III, *p* ≤ 0.02).

Thus, modality-specific semantic uncertainty did not have an invariant temporal neural signature. Its neural expression was shaped by the semantic-specification state of the accompanying modality, indicating that uncertainty defined at Level 1 was already expressed in a manner sensitive to the broader cross-modal configuration.

### Configuration-dependent neural expression of cross-modal uncertainty transformation

We next examined whether the bidirectional but asymmetric transformation identified computationally at Level 2 was likewise dynamically expressed across semantic configurations.

Gesture gain was associated with early N1-range activity when gesture remained pre-threshold (Types I and II, *p* ≤ 0.018, **Fig. 4a**), whereas speech gain showed broader P2- and P600-range associations when speech remained pre-threshold (Types I and III, *p* ≤ 0.022, **Fig. 4b**). When the two modalities differed in threshold status (Types II and III), both gesture and speech gain were associated with activity during the N1-P2 interval (*p* ≤ 0.011).

**Fig. 4.**
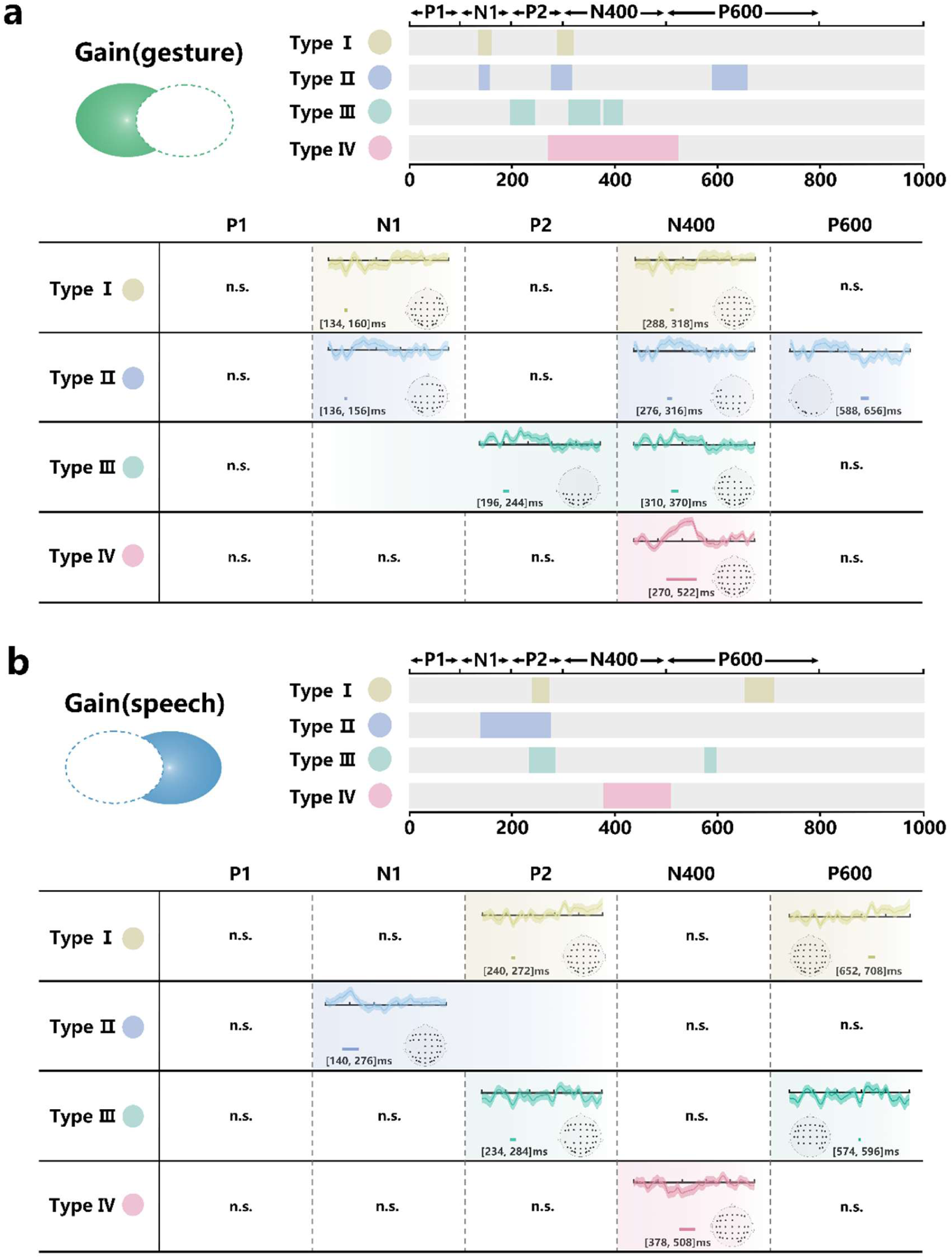
Cross-modal semantic configuration shapes the neural expression of cross-modal uncertainty transformation. **a,b**, Spatiotemporal associations between the semantic-congruency difference wave and gesture gain (**a**) or speech gain (**b**) across the four cross-modal semantic configurations. Gesture and speech gain quantified changes in modality-specific entropy from the unisensory to the corresponding multisensory context; positive values indicate reduced uncertainty, and negative values indicate increased uncertainty. Participant-level Spearman correlation coefficients were tested against zero using two-tailed spatiotemporal cluster-based permutation tests with 1,000 permutations. **a**, Gesture gain was associated with early N1 activity when gesture remained pre-threshold (Types I and II) and with N400 activity across all four configurations. **b**, Speech gain showed P2- and P600-range associations when speech remained pre-threshold (Types I and III), whereas an N400 association was most evident when both modalities were threshold-reached (Type IV). When the two modalities differed in threshold status (Types II and III), both gesture and speech gain were associated with activity in the N1-P2 interval.

At later latencies, gesture gain was associated with N400 activity across all four configurations (Types I-IV, *p* ≤ 0.016), whereas an N400 association with speech gain was observed primarily when both modalities were threshold-reached (Type IV, *p* = 0.002).

These results extend the computational asymmetry observed at Level 2 to its neural expression. Gesture- and speech-related uncertainty transformations showed different temporal profiles, and these profiles varied with the joint semantic specification of the two modalities. Cross-modal transformation was therefore not expressed as a uniform reduction signal, but as a configuration-dependent and modality-asymmetric component of the hierarchy.

### Configuration-dependent neural expression of multisensory uncertainty convergence

We finally examined whether multisensory convergence at Level 3 was similarly shaped by cross-modal semantic configuration. AFE showed robust associations with N400 activity across all four configurations (Types I-IV, *p* < 0.024, **Fig. 5a**). When both modalities remained pre-threshold (Type I), AFE-related activity additionally extended into the early N1 stage (*p* = 0.003), indicating that neural sensitivity to this fusion-based convergence was present even before either modality had reached its semantic-identification threshold.

**Fig. 5.**
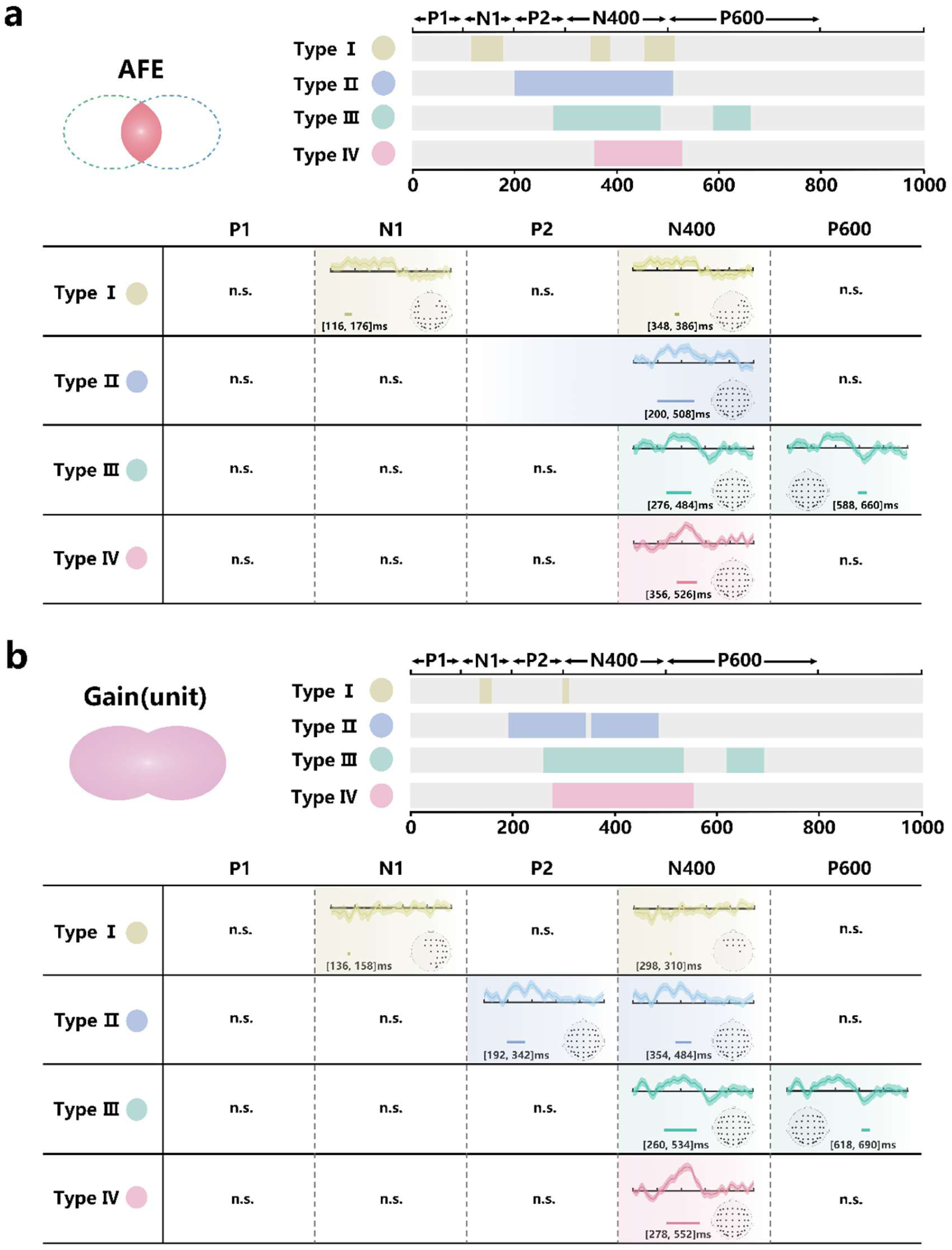
Cross-modal semantic configuration shapes the neural expression of multisensory uncertainty convergence. **a,b**, Spatiotemporal associations between the semantic-congruency difference wave and aligned fusion entropy (AFE; **a**) or unit gain (**b**) across the four cross-modal semantic configurations. AFE quantified the change in uncertainty associated with uncertainty-weighted fusion of the aligned gesture and speech semantic probability distributions, whereas unit gain quantified the change in uncertainty from the initial gesture representation to the subsequent speech representation in the multisensory context. Participant-level Spearman correlation coefficients were tested against zero using two-tailed spatiotemporal cluster-based permutation tests with 1,000 permutations. **a**, AFE was associated with N400 activity across all four configurations and additionally with early N1 activity when both modalities remained pre-threshold (Type I). **b**, Unit gain showed N400 associations when at least one modality was threshold-reached (Types II-IV); effects in Type I were weaker and more temporally restricted. Both AFE and unit gain additionally showed P600-range associations when gesture was threshold-reached and speech remained pre-threshold (Type III).

Unit gain showed a different temporal profile. N400 associations were observed when at least one modality was threshold-reached (Types II-IV, *p* < 0.001, **Fig. 5b**), with the largest observed *t* statistic in Type IV (t_max_ = 4.15). In Type I, when both modalities were pre-threshold, unit-gain-related effects were weak and transient (*p* < 0.05).

Both AFE and unit gain showed additional P600-range associations when gesture was threshold-reached while speech remained pre-threshold (Type III, *p* < 0.05, **Fig. 5**). This configuration also showed later associations with speech entropy and speech gain **(Fig. 3b and 4b),** indicating that multiple computational components remained neurally expressed at later latencies when speech semantic specification was incomplete.

Together, these findings show that multisensory convergence was not associated with a single invariant neural signature. Moreover, the distinct profiles of AFE and unit gain paralleled their computational dissociation, further indicating that convergence reflects complementary aspects of the integrated representation rather than a single endpoint of progressive uncertainty reduction.

Across all levels, cross-modal semantic configuration therefore shaped the neural expression of uncertainty-related quantities. The computational hierarchy was therefore not implemented as an invariant temporal cascade from Level 1 to Level 3. Instead, its component processes were dynamically reconfigured according to the semantic information available from gesture and speech.

### Statistical dependencies connect the levels of hierarchical uncertainty transformation

The preceding analyses showed that the three computational levels were distinguishable in both their computational profiles and neural expression, while their temporal organization varied with cross-modal semantic configuration. We next asked whether these levels were statistically coupled, as expected if they reflect interdependent components of a common uncertainty-transformation process rather than independent descriptive measures. We therefore used mediation and moderated mediation analyses to characterize the statistical dependencies among these computational measures and their relationships with neural and behavioral indices of semantic integration.

### Statistical dependency between complementary measures of multisensory convergence

We first examined the relationship between the two Level 3 convergence measures and their association with neural and behavioral indices of semantic integration. Unit gain significantly mediated the statistical association between AFE and the N400 semantic congruency effect (indirect β = 0.54, 95%CI = [0.26, 0.85]) **(Fig. 6b)**, while exerting a strong direct effect itself (β = 0.54, 95%CI = [0.26, 0.85]) **(Fig. S3b)**. Parallel analyses using the behavioral reaction-time congruency effect as the outcome showed a comparable mediation pattern (**Fig. 6c and Fig. S3d**). These findings indicate that AFE and unit gain, although computationally complementary, were not independent. Their statistical relationship was associated with variation in both neural and behavioral indices of multimodal semantic integration, linking the computational characterization of multisensory convergence to its experimentally observed consequences.

**Fig. 6.**
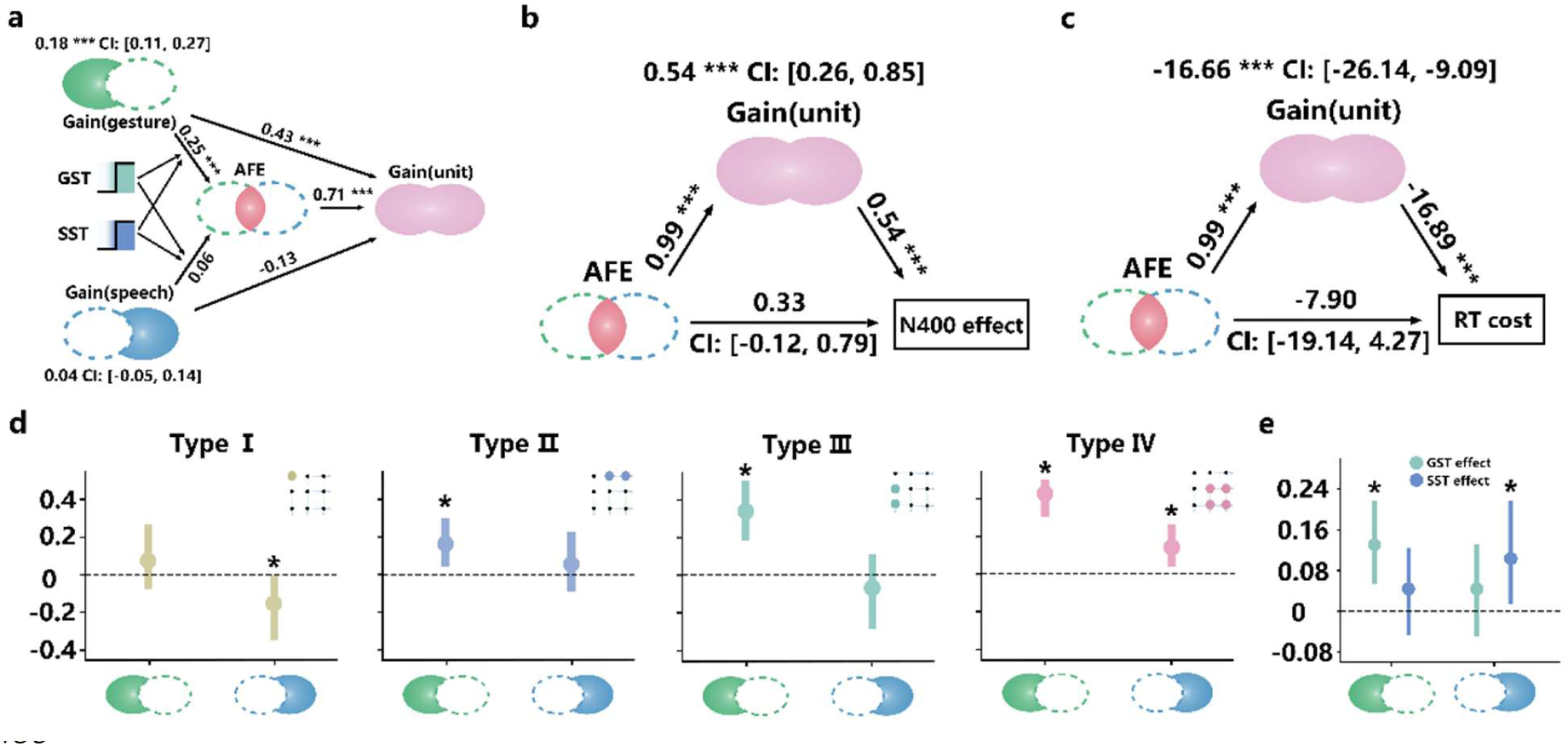
Configuration-dependent statistical dependencies across the uncertainty-transformation hierarchy. **a**, Cross-level mediation analysis relating modality-specific uncertainty transformation to multisensory convergence. Gesture and speech gains were entered as predictors of AFE, with unit gain as the outcome. Gesture gain showed both a direct association with unit gain and an indirect association through AFE, whereas no reliable overall effect was observed for speech gain. **b**, Mediation analysis relating the two multisensory convergence measures to the neural semantic-congruency effect. Unit gain mediated the statistical association between AFE and the N400 semantic-congruency effect. **c**, Corresponding mediation analysis using the behavioral reaction-time semantic-congruency effect as the outcome. **d**, Conditional indirect effects of gesture and speech gain on unit gain through AFE across the four cross-modal semantic configurations. Gesture-related indirect effects were significant when at least one modality had reached its semantic-identification threshold (Types II-IV). In contrast, the indirect effect of speech gain was significant only when gesture and speech shared the same threshold status: it was negative when neither modality had reached its threshold (Type I) and positive when both modalities had reached their respective thresholds (Type IV). **e**, Moderated mediation analyses testing whether the Gain(unisensory)-AFE-Gain(unit) pathway varied with the corresponding modality-specific semantic-specification state, coded as 0 for pre-threshold and 1 for threshold-reached. Significant indices of moderated mediation were observed for both gesture and speech modalities. Indirect effects were estimated using 5,000 bootstrap samples and 95% bias-corrected confidence intervals. Analyses were conducted across 171 stimulus-condition observations. Error bars indicate 95% confidence intervals.

We next asked whether this dependency varied with semantic specification. Across the four cross-modal semantic configurations **(Fig. S3e)**, unit gain mediated the association between AFE and the N400 congruency effect, with significant indirect effects across all configurations (β = 0.34 - 0.93, all 95%CIs excluded zero), and no reliable direct effect of AFE on the N400 response **(Fig. S3i)**. Crucially, the strength of this mediated pathway varied with gesture semantic specification (Index of moderated mediation = 0.24, 95%CI = [0.08, 0.50]), whereas moderation by speech semantic specification was not significant. Thus, even within the convergence level, statistical coupling of the two complementary measures varied, with gesture specification exerting a stronger influence on how AFE was related to neural semantic integration through unit gain.

### Cross-level dependencies link uncertainty transformation to multisensory convergence

We next tested the central cross-level prediction of the hierarchical account: whether modality-specific uncertainty transformation at Level 2 was statistically linked to multisensory convergence at Level 3. In a cross-level mediation model, gesture and speech gains were entered as modality-specific indices of uncertainty transformation, AFE as the mediator and unit gain as the convergence outcome.

Gesture gain showed both a significant direct association with unit gain (β = 0.43, 95%CI = [0.31, 0.54]) and an indirect association through AFE (β = 0.18, 95%CI = [0.11, 0.27]), whereas no reliable overall effect was observed for speech gain **(Fig. 6a)**. Thus, cross-modal changes in gesture uncertainty were statistically linked to multisensory convergence both directly and through the fusion-based convergence measure. More generally, these findings connect Level 2 transformation to Level 3 convergence, supporting the view that the computational levels describe interdependent components of a common uncertainty-transformation process rather than independent metrics.

Critically, however, the absence of an overall speech-gain effect did not imply that speech-related uncertainty transformation was unrelated to convergence. Rather, moderated mediation analyses showed significant moderation of the indirect pathways for both gesture gain (Index of moderated mediation = 0.13, 95%CI = [0.06, 0.21]) and speech gain (Index = 0.10, 95%CI = [0.02, 0.21]) **(Fig. 6e)**. Conditional analyses revealed distinct configuration-dependent profiles (**Fig. 6d**). The indirect association between gesture gain and unit gain through AFE was significant when one or both modalities had reached the semantic-identification threshold (Type II: 0.16, 95%CI = [0.05, 0.30]; Type III: 0.34, 95%CI = [0.18, 0.54], *p* < 0.001; Type IV: 0.43, 95%CI = [0.30, 0.57]) **(Fig. 6d)**. In contrast, the speech-gain pathway showed a more restricted configuration-dependent pattern. The estimated indirect effect was negative when both modalities remained pre-threshold (Type I: 95%CI = [-0.30, -0.01]), whereas a positive indirect effect was observed when both modalities were threshold-reached (Type IV: 95%CI = [0.04, 0.26]). Thus, the statistical coupling between Level 2 transformation and Level 3 convergence did not follow a uniform pattern across modalities or semantic configurations.

Together, these mediation analyses provide a complementary form of evidence for the hierarchical account. The computational levels were distinguishable in their definitions, profiles and neural associations, yet remained statistically interconnected. Moreover, the strength and pattern of these cross-level dependencies varied with cross-modal semantic configuration. Hierarchical uncertainty transformation therefore reflects neither three independent quantities nor a fixed serial cascade, but a set of computationally differentiated and statistically coupled processes whose organization depends on the semantic information available from each modality.

### Integrated summary

Across computational, neural and cross-level analyses, multimodal semantic integration was characterized by a structured hierarchy of uncertainty transformation rather than by a uniform reduction of uncertainty (**Fig. 7**). At Level 1, semantic uncertainty varied systematically within modality-specific gesture and speech representations, establishing the uncertainty structure available for cross-modal transformation. At Level 2, cross-modal information produced bidirectional but asymmetric changes in modality-specific uncertainty, inconsistent with a simple uniform-reduction account. At Level 3, AFE and unit gain showed distinct computational profiles, indicating that multisensory convergence extended beyond modality-specific uncertainty changes and comprised complementary aspects of uncertainty at the integrated level.

**Fig. 7.**
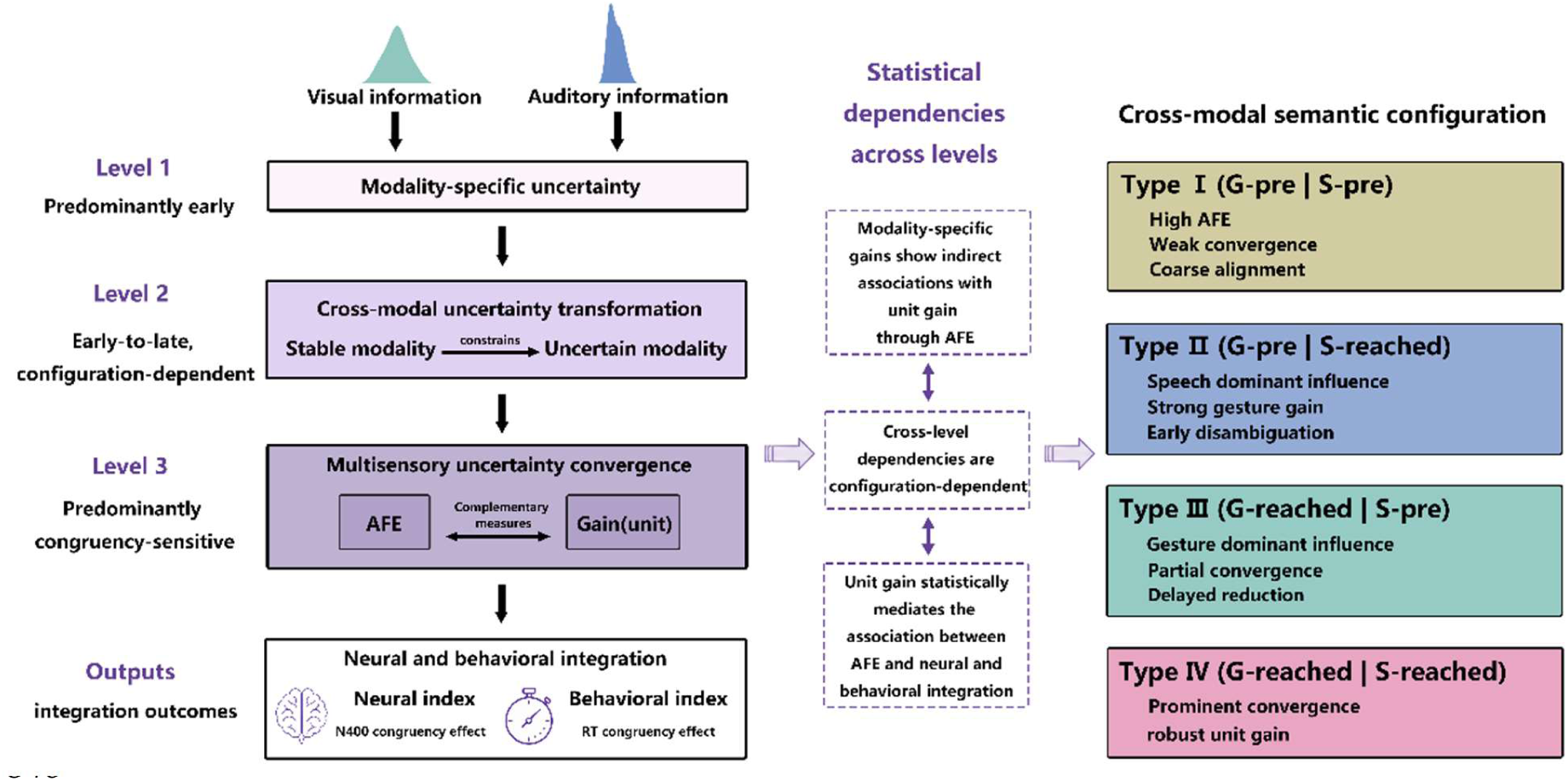
Integrated framework for hierarchical uncertainty transformation during multimodal semantic integration. The framework summarizes the computational, neural and statistical organization of hierarchical uncertainty transformation during gesture-speech integration. **Left**, uncertainty changes computational form across three levels: modality-specific uncertainty (Level 1), cross-modal uncertainty transformation (Level 2) and multisensory uncertainty convergence (Level 3), culminating in neural and behavioral indices of semantic integration. Level 3 is characterized by complementary convergence measures aligned fusion entropy (AFE) and unit gain. **Middle**, statistical dependencies link the computational levels: modality-specific gains show indirect associations with unit gain through AFE, and unit gain statistically mediates the association between AFE and neural and behavioral integration. These cross-level dependencies vary with semantic specification. **Right**, the four cross-modal semantic configurations, defined by the joint threshold status of gesture and speech, differentially shape the neural expression and statistical coupling of the hierarchy.

The neural results provided a second line of support for this computational hierarchy. Modality-specific uncertainty was preferentially associated with modality-related neural responses, whereas cross-modal uncertainty transformation and multisensory convergence were more closely associated with semantic-congruency-sensitive activity. Configuration-resolved analyses further revealed distinct temporal profiles across the hierarchy: modality-specific uncertainty showed early associations across configurations but configuration-dependent later effects; cross-modal transformation showed asymmetric gesture- and speech-related profiles; and multisensory convergence was consistently associated with N400 activity, with unit-gain-related expression becoming more prominent once at least one modality reached its semantic-identification threshold. Thus, the computational hierarchy was neurally differentiated but temporally overlapping, with its expression dynamically shaped by the joint semantic specification of gesture and speech rather than organized as a fixed sequence of discrete neural stages.

Finally, mediation analyses showed that computational differentiation did not imply independence. Cross-modal uncertainty transformation at Level 2 was statistically linked to multisensory convergence at Level 3, and these cross-level dependencies varied with the joint semantic specification of gesture and speech. Thus, uncertainty did not simply decrease as multimodal semantic information accumulated. Rather, it changed computational form—from modality-specific uncertainty, through asymmetric cross-modal transformation, to multisensory convergence—across distinguishable but interdependent levels whose neural expression and statistical coupling were dynamically shaped by cross-modal semantic configuration.

## Discussion

The present study supports a hierarchical uncertainty transformation account of multimodal semantic integration. Across computational analyses, modality-specific semantic uncertainty, cross-modal uncertainty transformation and multisensory uncertainty convergence showed distinct profiles. These computational distinctions were accompanied by differentiated but temporally overlapping neural associations, whose expression varied systematically with cross-modal semantic configuration. Cross-level analyses further revealed statistical dependencies between cross-modal transformation and multisensory convergence, with these dependencies themselves varying across configurations. Together, these findings indicate that multimodal integration is not adequately characterized by a uniform reduction of uncertainty as information accumulates. Rather, uncertainty is reorganized as modality-specific semantic representations interact and converge.

The central implication is that uncertainty need not characterize the same computational property throughout multimodal integration. Within modality-specific representations, uncertainty reflects the distribution of competing semantic interpretations; during cross-modal interaction, it reflects changes in those modality-specific distributions; and at multisensory convergence, it characterizes uncertainty associated with the convergence of modality-specific semantic information. Hierarchical uncertainty transformation therefore refers to a change in what uncertainty characterizes, rather than to the progressive reduction of a single uncertainty signal through successive processing stages. This distinction complements existing probabilistic accounts of multisensory processing. Bayesian and predictive-processing frameworks provide principled accounts of evidence weighting and representational updating over probability distributions^30–33^. The present framework highlights an additional organizational property that may be particularly relevant to the hierarchically organized multisensory system^24,34–36^: changes in uncertainty within one computational level need not be accompanied by equivalent changes at another. The distinct profiles of modality-specific entropy, cross-modal gains and convergence measures are consistent with this view. The proposed hierarchy should therefore be understood neither as an alternative to probabilistic inference nor as a fixed serial sequence, but as a computational description of how distinct forms of uncertainty are related as distributed semantic representations interact and converge.

A second contribution is an explicit computational procedure for aligning and combining heterogeneous semantic distributions. Gesture and speech naming distributions were first mapped onto a shared probabilistic semantic space, such that corresponding semantic alternatives occupied common dimensions while modality-specific alternatives were retained. This operation is conceptually compatible with computational accounts in which modality-specific information converges within a shared multimodal semantic system^37,38^. The aligned distributions were then combined using uncertainty-dependent modality weights derived from their entropy, such that distributions concentrated over fewer semantic alternatives contributed more strongly to the fused distribution. In this formulation, uncertainty is not only a property of the representations being integrated, but also a computational variable regulating their relative contribution to fusion. This differs in emphasis from conventional reliability- or precision-weighting formulations^15,39^ because the relevant uncertainty is defined over competing semantic interpretations rather than sensory noise or measurement variance. Importantly, entropy-dependent weighting is not inherently incompatible with Bayesian inference; instead, it provides a complementary operationalization for situations in which the uncertainty relevant to integration resides in the structure of semantic probability distributions. The combination of representational alignment and uncertainty-dependent fusion thus provides a tractable way to integrate modality-specific semantic distributions that initially differ in their sets of semantic alternatives.

The neural results provided complementary evidence for the temporal organization of hierarchical uncertainty transformation. The overall neural profile suggested an early emergence of gesture-related semantic uncertainty, followed by speech-related semantic processing, whereas bidirectional cross-modal transformation and multisensory convergence were expressed most prominently in the N400 range, which is broadly sensitive to semantic integration or facilitated lexical access^40,41^. This pattern suggests a broad temporal organization in which modality-related semantic information becomes available before uncertainty transformation and convergence are prominently expressed during cross-modal semantic interaction. Configuration-resolved analyses further showed that this organization was dynamically modulated by the semantic information available from the two modalities. Early modality-specific associations were relatively consistent across configurations, whereas later associations became increasingly configuration-dependent; cross-modal transformation showed asymmetric gesture- and speech-related profiles, indicating that the neural consequences of uncertainty in one modality depended on the semantic information available from the other^42,43^. Convergence-related activity was prominent in the N400 range across configurations, with unit-gain-related associations becoming more evident once semantic information was specified in one or both modalities. Thus, the computational hierarchy exhibited a broad temporal organization but did not map onto a fixed sequence of discrete neural stages. Instead, its temporal expression was dynamically reconfigured according to the joint semantic specification of gesture and speech.

The mediation analyses extended this configuration dependence from neural expression to statistical relationships among computational quantities. Within multisensory convergence, unit gain statistically mediated the association between aligned fusion entropy (AFE) and neural and behavioral indices of semantic integration, indicating that the two convergence measures were distinguishable yet interconnected. Across levels, gesture gain showed both a direct association with unit gain and an indirect association through AFE, whereas no reliable overall effect was observed for speech gain. These dependencies further varied with semantic specification. The indirect gesture-gain pathway was evident when one or both modalities had reached their semantic-identification thresholds and was numerically larger when gesture itself was threshold-reached, whereas the speech-gain pathway showed a more restricted configuration-dependent profile. Thus, computational differentiation did not imply independence: the statistical relationships between cross-modal uncertainty transformation and multisensory convergence changed with the semantic information available from the two modalities. These analyses do not establish a causal sequence among computational levels but provide complementary evidence that the proposed hierarchy comprises statistically interconnected rather than isolated computational quantities.

Further insight into multisensory convergence comes from the dissociation between AFE and unit gain. Although both characterize convergence at level 3, they quantify different properties of the integrated computational context. AFE relates modality-specific uncertainty to the entropy of the uncertainty-weighted fused semantic distribution, whereas unit gain quantifies the change in uncertainty relative to the initial gesture representation. Their distinct computational profiles and neural associations therefore indicate that multisensory convergence cannot be reduced to a single measure of uncertainty reduction. Instead, convergence encompasses complementary information-theoretic properties that become differently expressed as modality-specific information is combined. This distinction further supports the view that hierarchical uncertainty transformation reflects a structured reorganization of uncertainty rather than a unitary reduction process^44,45^.

More broadly, the present framework suggests that the organization of uncertainty may be important whenever coherent representations must be constructed from heterogeneous information sources. When multiple representational streams support partially different alternatives and evolve over different timescales, the computational problem may not be exhausted by asking how uncertainty decreases as additional evidence becomes available. It may also be necessary to consider how uncertainty-bearing representations are coordinated as information from different sources is brought into relation. Gesture-speech communication provides a useful test case because the two modalities differ in representational format and temporal structure while contributing to a common semantic interpretation. Related computational problems arise in audiovisual speech^46,47^ and other forms of naturalistic multimodal communication^48,49^. Whether the same hierarchical organization extends to these domains remains an empirical question. The present findings therefore motivate hierarchical uncertainty transformation as a candidate principle for understanding distributed multimodal integration, rather than establishing it as a universal property of cognitive systems.

Several limitations define the scope of this account. First, the semantic probability distributions were derived from discrete naming responses. This provides an interpretable probability space over competing meanings but does not capture the richer geometry of continuous semantic representations; extending the framework to embedding-based or other continuous semantic spaces will therefore be important^21,50–53^. Second, EEG provides the temporal resolution required to characterize overlapping neural associations but offers limited anatomical precision. Combining the present framework with spatially resolved neural measurements could determine how these computational quantities are distributed across cortical systems^54^. Finally, the mediation analyses characterize statistical dependencies among stimulus-condition-level quantities and cannot establish causal relationships among computational levels. Experimental manipulation of specific computational quantities will be required to determine whether the observed dependencies reflect directional mechanisms^55,56^. In conclusion, the present findings support a hierarchical uncertainty transformation account of multimodal semantic integration, in which uncertainty changes computational form as modality-specific semantic representations interact and converge. By integrating heterogeneous semantic distributions within a common information-theoretic framework, this account shifts the computational question from how much uncertainty is reduced to how uncertainty is reorganized across interacting representations. This perspective provides a new way to understand how coherent meaning emerges from distributed and uncertain information.

## Methods

### Participants

A total of 120 native Chinese speakers participated in four experiments (30 participants per experiment). **Experiments 1-2** quantified semantic uncertainty associated with speech and gesture, respectively (**Experiment 1**: 17 females, age = 24 ± 5.41 years; **Experiment 2:** 15 females, age = 23 ± 4.90 years). **Experiment 3** quantified semantic uncertainty during multisensory gesture-speech processing (15 females, age = 22 ± 2.66 years), and **Experiment 4** examined the neural dynamics of gesture-speech integration using EEG (16 females, age = 22 ± 3.43 years). All participants were right-handed (Edinburgh Handedness Inventory laterality quotient^57^ = 89.20 ± 14.69), reported normal hearing and had normal or corrected-to-normal vision. Participants provided informed consent and received ¥100 per hour as compensation. The study was approved by the Ethics Committee of the Institute of Psychology, Chinese Academy of Sciences (no. H21077).

### Stimuli

The stimulus set consisted of 19 gestures and 19 corresponding speech recordings collected previously from two native Chinese speakers (one male, one female)^58,59^ (**Table S1**). Two orthogonal factors were manipulated: gender congruency (same-versus opposite-gender voice relative to the gesturer) and semantic congruency (congruent versus incongruent gesture-speech pair).

Four independent pre-tests were conducted to validate the stimuli and determine the temporal points at which semantic information became identifiable within each modality. In **pre-test 1** (n = 32, 16 females, age = 22 ± 4.48 years), participants rated the semantic relatedness of gesture-speech pairs on a five-point scale (1 = no relation, 5 = very strong relation), confirming high ratings for congruent pairs (Mean = 4.47, SD = 0.42) and low ratings for incongruent pairs (Mean = 1.44, SD = 0.43). **Pre-test 2** (n = 30, 17 females, age = 24 ± 4.56 years) provided an independent behavioral validation of the semantic congruency manipulation, with longer reaction times for incongruent pairs (Mean = 556.47 ± 11.68 ms) than congruent pairs (Mean = 530.85 ± 12.33 ms; t(29) = 7.21, *p* < 0.001).

**Pre-tests 3** (n = 30, 16 females, age = 24 ± 4.37 years) and **4** (n = 30, 15 females, age = 22 ± 3.26 years) determined the DP for gestures and IP for speech, respectively. Participants viewed or heard stimulus fragments of progressively increasing duration (starting at 40 ms) and produced a single verb describing each fragment. The DP or IP was defined as the earliest fragment duration at which the same semantic response remained stable across four to six consecutive presentations. The mean DP for gestures was 184.82 ms (SD = 83.79) and the mean IP for speech was 177.54 ms (SD = 67.69) (**Table S2**).

Gesture videos had a mean duration of 1,771 ms (SD = 308), and speech recordings had a mean duration of 447 ms (SD = 93). For each stimulus, three fragments were generated relative to its modality-specific semantic identification threshold: a pre-threshold fragment (0.75 × DP/IP), a threshold fragment (1 × DP/IP) and a post-threshold fragment (1.25 × DP/IP). These were denoted BD, DP and AD for gesture and BI, IP and AI for speech, respectively. Crossing the three speech and the three gesture semantic specification states yielded nine fragment conditions.

Fragment duration therefore manipulated semantic specification, whereas semantic uncertainty was quantified independently from the empirical probability distribution of naming responses using Shannon entropy (**Fig. S4**).

### Cross-modal semantic configurations

To examine whether multimodal integration differed according to the joint semantic specification of the two modalities, the three semantic-specification states within each modality were further classified according to whether the independently determined semantic-identification threshold had been reached. For gesture, DP and AD fragments were classified as threshold-reached, whereas BD fragments were classified as pre-threshold. For speech, IP and AI fragments were classified as threshold-reached, whereas BI fragments were classified as pre-threshold.

Crossing these binary semantic-specification states yielded four cross-modal semantic configurations: Type I, gesture pre-threshold & speech pre-threshold; Type II, gesture pre-threshold & speech threshold-reached; Type III, gesture threshold-reached & speech pre-threshold; and Type IV, gesture threshold-reached & speech threshold-reached (**Fig. 1a**).

The four configurations did not discretize semantic uncertainty. All stimulus-level information-theoretic measures were retained as continuous variables within each configuration. The original three-level semantic-specification manipulation (BD/DP/AD × BI/IP/AI) was retained for analyses requiring the full factorial structure.

### Unisensory naming experiments

**Experiments 1 and 2** were used to estimate stimulus-specific semantic response distributions for speech and gesture, respectively. Participants heard isolated speech fragments or viewed isolated gesture fragments in randomized order and typed a verb corresponding to the perceived meaning. Each fragment was presented twice for both the male and female performers to balance gender effects, resulting in 114 trials per participant.

For each stimulus and fragment condition, naming responses were aggregated across participants. The frequency of each unique semantic response was divided by the total number of valid responses to obtain an empirical probability distribution over semantic labels. These distributions were used to calculate stimulus-specific unisensory semantic uncertainty.

### Multisensory naming experiment

**Experiment 3** was used to estimate semantic response distributions when each target modality was processed in the presence of the other modality. Participants completed two task phases using the 171 semantically congruent gesture-speech combinations generated by crossing 19 stimulus identities with the nine fragment combinations. Stimuli were distributed across nine blocks. Six semantically incongruent pairs were additionally inserted into each block to reduce the predictability of semantic congruency.

Each stimulus was presented twice across the gender-congruency manipulation to balance gender effects. In one phase, participants named the gesture while ignoring the accompanying speech; in the other phase, they named the speech while ignoring the gesture. The order of the two task phases was counterbalanced across participants. Blocks containing shorter fragments preceded those containing longer fragments to minimize carry-over of semantic information from more identifiable to less identifiable fragments. Each participant completed 450 trials in approximately 1 hour.

Naming responses were aggregated separately for each stimulus, fragment condition and target modality and normalized to yield multisensory gesture and speech probability distributions. These distributions were used to calculate stimulus-specific multisensory semantic uncertainty.

### EEG experiment

**Experiment 4** used a semantic priming paradigm in which a gesture fragment served as the prime and was followed immediately by speech^60^. Semantic congruency was manipulated between gesture and speech, whereas gender congruency served as an orthogonal, task-irrelevant factor.

Each trial began with a fixation cross presented for 0.5-1.5s, followed by the gesture fragment and then the speech stimulus. Participants performed a speeded two-alternative forced-choice judgement of speaker gender using left- and right-button presses. The orthogonal gender-classification task minimized explicit semantic decision demands while allowing semantic congruency effects to index implicit gesture-speech semantic integration^59,60^. Participants completed 1,368 trials in a single session. Stimuli were presented using Presentation software (Neurobehavioral Systems).

### Construction of a shared probabilistic semantic space

Gesture and speech naming responses were defined over partially overlapping semantic vocabularies. To enable direct comparison of the two modality-specific probabilistic distributions, responses were projected into a shared probabilistic semantic space with a common set of semantic dimensions.

The shared space was constructed by taking the union of all semantic labels produced in the gesture and speech naming experiments. Semantically identical labels across modalities were aligned to the same dimension, whereas labels occurring in only one modality were retained as modality-specific dimensions. For each modality-specific dimension, the corresponding probability in the other modality was set to zero. This alignment yielded a common set of semantic dimensions while preserving all observed responses and their modality-specific probabilities.

For each stimulus and condition, response frequencies were normalized within each modality and projected onto the aligned semantic dimensions, yielding a gesture probability vector, *P*_(G)_, and a speech probability vector, *P*_(S_). Their corresponding elements, *P*_(G)_(*i*) and *P*_(S)_(*i*), denote the probabilities assigned by gesture and speech, respectively, to the same semantic label. The resulting vectors therefore had identical dimensionality and a one-to-one correspondence between semantic dimensions, providing the common probabilistic space used for subsequent uncertainty-weighted semantic fusion.

### Uncertainty-weighted semantic fusion

Following alignment in the shared probabilistic semantic space, the gesture and speech probability distributions, *P*_(G)_ and *P*_(S)_, were combined according to their semantic uncertainty. Uncertainty was quantified by the entropy of each distribution (*H*(*G*) and *H*(*S*); see *Hierarchical information-theoretic metrics below*) and converted to positive weighting quantities as 2^−H(G)^ and 2^−H(S)^.

The resulting quantities were normalized across modalities to obtain the fusion weights:

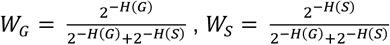

For each semantic dimension *i* in the shared space, the fused probability *P*_(GS)_ was computed as:

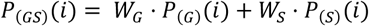

Where *P*_(G)_(*i*) and *P*_(S)_(*i*) denote the normalized probabilities assigned to semantic label (*i*) in the aligned gesture and speech distributions, respectively. Thus, the relative contribution of each modality varied with the uncertainty of its semantic distribution, with lower-entropy distributions contributing proportionally more to the fused representation.

The entropy of the resulting fused distribution, *H*_(GS)_, was calculated using the same Shannon entropy formulation and quantified the residual uncertainty of the uncertainty-weighted multimodal representation. The aligned modality-specific distributions and the resulting fused distribution provided the quantities required for the multisensory convergence measures described below.

### Hierarchical information-theoretic metrics

We characterized the hierarchical transformation of semantic uncertainty during multimodal integration using information-theoretic measures at three computational levels: modality-specific semantic uncertainty, cross-modal uncertainty transformation and multisensory uncertainty convergence. These levels capture complementary aspects of the integration process, from uncertainty within individual modality-specific representations, through changes in that uncertainty in the presence of cross-modal information, to its reorganization as gesture and speech converge into a multimodal representation. The levels were defined as distinct computational quantities rather than discrete temporal processing stages; their temporal neural expression was subsequently examined using EEG.

#### *Level 1*: Modality-specific semantic uncertainty

At the first level, Shannon entropy was used to quantify uncertainty within the modality-specific semantic representations. For a discrete probability distribution *P* over semantic labels, entropy was defined as:

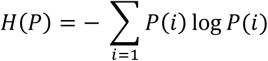

where *P*(*i*) denotes the probability assigned to semantic label (*i*). Lower entropy indicates a more concentrated response distribution, with probability mass concentrated on fewer semantic alternatives, whereas higher entropy reflects a more dispersed distribution across alternative semantic interpretations.

For each stimulus and fragment condition, entropy was calculated separately from the gesture and speech distributions obtained in the unisensory and multisensory naming experiments. This yielded four modality-specific measures:

1. Unisensory speech entropy: H(*unispeec*ℎ);
2. Multisensory speech entropy: H(*multispeec*ℎ);
3. Unisensory gesture entropy: H(*unigesture*);
4. Multisensory gesture entropy: H(*multigesture*).

Entropy values were approximately normally distributed across conditions (**Fig. S4**).

#### *Level 2*: Cross-modal uncertainty transformation

At the second computational level, we quantified changes in modality-specific semantic uncertainty when information from the other modality was available. For each stimulus and condition, the entropy of the modality-specific distribution obtained in the multisensory context was compared with that obtained in the corresponding unisensory context.

Gesture gain was defined as: Gain(*gesture*) = H(*unigesture*) − H(*multigesture*); and speech gain as: Gain(*speec*ℎ) = H(*unispeec*ℎ) − H(*multispeec*ℎ).

Positive values indicate reduced uncertainty in the multisensory relative to the corresponding unisensory context, and values close to zero indicate little change. The gain measures therefore capture the magnitude of cross-modal uncertainty transformation.

#### *Level 3*: Multisensory uncertainty convergence

At the third computational level, we quantified how modality-specific semantic uncertainty was reorganized as gesture and speech converged into a multimodal representation. Whereas the preceding level characterized cross-modal transformations of gesture and speech uncertainty separately, this level characterized the resulting organization of uncertainty across modalities.

Two complementary measures were used: AFE and unit gain.

AFE was defined as:

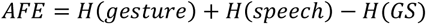

where *H*(*gesture*) and *H*(*speech*) denote the entropy of the unisensory gesture and speech probability distributions, respectively, and *H*(*GS*) denotes the entropy of the uncertainty-weighted fused distribution. AFE therefore indexes the change in uncertainty associated with combining the two modality-specific distributions within the shared probabilistic semantic space. Larger AFE values indicate a larger difference between the summed modality-specific uncertainty and the uncertainty of the fused representation, providing a fusion-based measure of multisensory convergence.

Unit gain was defined as:

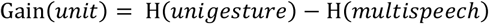

where H(*unigesture*) denotes semantic uncertainty in the unisensory gesture representation and H(*multispeec*ℎ) denotes uncertainty in the speech representation when speech was processed in the presence of the preceding gesture. Gesture served as the reference because it preceded speech as the semantic prime in the experimental sequence. Unit gain therefore quantified the cross-modal change in uncertainty from the initial gesture representation to the subsequent multisensory speech representation.

All information-theoretic measures were computed separately for each stimulus and condition and retained as continuous variables in subsequent analyses. Computational details and stimulus-level values are provided in Supplementary Material **Table S3 and S4**.

### EEG recordings and pre-processing

EEG data were recorded from 42 Ag/AgCl electrodes (10-20 system^61^) using a PORTI-32/MREFA amplifier (TMS International B.V., Enschede, NL) at 500 Hz (0.01 to 70 Hz bandpass). Vertical and horizontal eye movements were monitored with four electrodes placed above and below the left eye and at the bilateral outer canthi. Signals were online-referenced to the left mastoid and offline re-referenced to the average of both mastoids (impedance < 5 KΩ). Pre-processing in EEGLAB^62^ included 0.05 Hz high-pass and 30 Hz low-pass filtering, independent component analysis (ICA) based artifact removal (eye blinks and muscle activity), and epoching from -200ms to 1000ms relative to speech onset, with 200ms pre-stimulus baseline correction. Incorrect trials (11.49%) and participants with > 30% artifact rejection were excluded. Outliers were handled using a 2-SD trimmed mean procedure within each condition and participant.

### Time-resolved full-factorial RSA

To characterize the temporal organization of gesture-, speech- and integration-related neural representations, we performed time-resolved full-factorial RSA^63^ using the Python packages rsatoolbox and NeuroRA^64^.

For each participant and time point, 18 × 18 neural representational dissimilarity matrices (RDMs) were constructed from multivariate EEG activity across 42 electrodes. The 18 conditions comprised the factorial combination of three gesture semantic-specification states (BD, DP and AD), three speech semantic-specification states (BI, IP and AI), and two levels of semantic congruency (congruent and incongruent). Condition-specific EEG activity patterns were extracted at each time point, and pairwise Euclidean distances between conditions were used to construct the neural RDMs.

Three hypothesis-driven model RDMs represented gesture semantic specification, speech semantic specification and semantic congruency (**Fig. 2a**). The gesture and speech model RDMs encoded ordinal distances among their respective semantic specification states, with BD/BI, DP/IP and AD/AI assigned values of 0, 1 and 2, respectively. The semantic congruency model RDM encoded whether pairs of conditions belonged to the semantically congruent or incongruent categories (0 = congruent, 1 = incongruent). For example, comparison of the BD-AI-congruent and AD-IP-congruent conditions yielded predicted dissimilarities of 2, 1 and 0 in the gesture, speech and congruency model RDMs, respectively.

At each time point, the vectorized neural RDM was simultaneously regressed onto the three model RDMs. The resulting regression coefficients indexed the unique correspondence between neural representational geometry and each model RDM.

Group-level inference was performed using one-sample t-tests on participant-level regression coefficients. Multiple comparisons across time were controlled using cluster-based permutation testing (1,000 iterations). Candidate clusters comprised consecutive time points exceeding a threshold of *p* < 0.05, with a minimum duration of 10 ms. Cluster-level significance was assessed against a null distribution generated by random sign-flipping of participant-level regression coefficients.

### Computational-EEG correspondence analyses

We examined the relationship between the information-theoretic measures and EEG activity in two complementary analyses. First, RSA-guided correlation analyses tested whether measures at each computational level were associated with the corresponding neural representations identified by the full-factorial RSA. Second, configuration-resolved analyses mapped the spatiotemporal expression of these computational-neural relationships across the four cross-modal semantic configurations.

### RSA-guided computational-EEG correlations

For each representational component identified by the full-factorial RSA, ERP amplitudes were extracted from the corresponding significant EEG cluster and related to the stimulus-level information-theoretic measures (**Fig. 2b**).

For modality-specific semantic uncertainty (Level 1), gesture and speech entropy were correlated with ERP amplitudes extracted from the RSA-defined clusters associated with gesture and speech semantic specification, respectively. For cross-modal uncertainty transformation (Level 2) and multisensory uncertainty convergence (Level 3), the semantic congruency difference wave was used as the neural index of multisensory integration^3,65,66^. Difference-wave amplitudes were calculated between congruent and incongruent conditions within the RSA-defined cluster associated with semantic congruency and correlated with gesture gain, speech gain, AFE and unit gain.

Spearman correlations were computed across stimulus-condition observations (n = 171) between each information-theoretic measure and its corresponding EEG amplitude. Multiple comparison was controlled using false discovery rate (FDR) correction.

### Configuration-resolved spatiotemporal correlations

We next examined whether the neural expression of the three computational levels varied across the four cross-modal semantic configurations defined above (**Figs. 3-5**). In contrast to the RSA-guided analyses, which were restricted to RSA-defined clusters, these analyses were performed across EEG time points and electrodes separately within each configuration.

For each cross-modal semantic configuration, stimulus-level information-theoretic metrics were correlated with EEG activity at each time point and electrode. Consistent with the RSA-guided analyses, modality-specific gesture and speech entropy (Level 1) were correlated with ERP amplitudes, whereas gesture and speech gain (Level 2), AFE and unit gain (Level 3) were correlated with semantic-congruency difference-wave amplitudes. Spearman correlation coefficients were calculated separately for each participant, computational measure and configuration.

Group-level inference was performed on participant-level correlation coefficients, which were transformed to t-statistics at each time point and electrode. Multiple comparisons across time and electrodes were controlled using spatiotemporal cluster-based permutation testing implemented in the MNE-Python toolbox (1,000 permutations). Clusters were defined as contiguous time-electrode samples exceeding a threshold of *p* < 0.05. Statistical significance was determined against a null distribution generated via random sign-flipping.

### Mediation and moderated mediation analysis

Mediation and moderated mediation analyses were conducted using the R package *lavaan*^67^ to examine the statistical dependencies among information-theoretic measures across the proposed computational hierarchy and their relationships with neural and behavioral outcomes. These analyses tested whether modality-specific uncertainty, cross-modal uncertainty transformation and multisensory uncertainty convergence formed an interdependent computational structure, and whether these dependencies varied with the semantic-information configuration of the two modalities.

We first examined dependencies between the two multisensory convergence measures and their associations with neural and behavioral outcomes. In the primary model, AFE was specified as the predictor, unit gain as the mediator and the N400 semantic congruency effect, averaged over the classical ROI (300-500 ms, midline region)3 as the neural outcome (**Fig. 6b**). A complementary model reversed the roles of AFE and unit gain to evaluate the alternative dependency structure between the two convergence metrics (**Fig. S3b**). Parallel models were fitted using the behavioral reaction-time congruency effect as the output (**Fig. 6c** and **Fig. S3d**). Next, we examined dependencies across computational levels. Gesture and speech gains, which quantify the transformation of modality-specific uncertainty between unisensory and multisensory contexts, were specified as predictors, with AFE as the mediator and unit gain as the dependent variable (**Fig. 6a**). This model tested whether cross-modal transformations of modality-specific uncertainty were statistically linked to subsequent multisensory convergence through dependencies among the corresponding computational measures.

Finally, moderated mediation analyses tested whether these cross-level dependencies varied across cross-modal semantic configurations. The four configurations were defined by the combination of binary semantic specification states of gesture and speech (pre-threshold versus threshold-reached). In the moderated mediation models, the corresponding modality-specific semantic specification state (0 = pre-threshold, 1 = threshold-reached) was included as a moderator of the pathway from gesture or speech gains to AFE (**Fig. 6d-e)**.

All indirect effects were evaluated using 5,000 bootstrap resamples and were considered statistically significant when the 95% bias-corrected confidence intervals excluded zero. Models were fitted to 171 stimulus-condition observations. The mediation models were used to characterize statistical dependencies consistent with the proposed hierarchical uncertainty-transformation framework and were not interpreted as evidence of causal relationships.

### Visualization of time-resolved EEG results

For visualization, time-resolved RSA coefficients and correlation coefficients were standardized across time within each model or computational measure and baseline-corrected relative to the pre-stimulus interval (−200 to 0 ms). Temporal smoothing was applied only to displayed time courses; all statistical analyses were performed on unsmoothed data.

## Declarations

### Availability statement

Data supporting the main findings of the study, along with the code illustrating key analyses, are available at https://osf.io/3yw4h/files/osfstorage.

### Competing interests

The authors declare the following competing interests: W. Y. Z. is an inventor on granted patent CN Patent No. ZL202522254705.3 entitled “A Brain-Inspired Approach and System for Probabilistic Alignment and Integration Measurement of Multimodal Semantic Representations”, granted on 17 February 2026. This patent covers the core algorithm for probabilistic representation alignment in gesture-speech integration described in this manuscript. The remaining authors declare no competing interests.

### Funding

This research was supported by grants from the STI 2030-Major Projects (2021ZD0201500) and Beijing Natural Science Foundation (4262073).

### Author contributions

Conceptualization, W.Y.Z. and Y.D.; Investigation, W.Y.Z., J.W.Y., and Z.Y.L.; Formal Analysis, W.Y.Z., J.W.Y., and Z.Y.L.; Methodology, W.Y.Z., J.W.Y., and Z.Y.L.; Validation, J.W.Y., and Z.Y.L.; Visualization, W.Y.Z., J.W.Y., and Z.Y.L.; Funding Acquisition, W.Y.Z. and Y.D.; Supervision, Y.D.; Project administration, Y.D.; Writing - Original Draft, W.Y.Z., J.W.Y., and Z.Y.L.; Writing - Review & Editing, W.Y.Z., J.W.Y., Z.Y.L. and Y.D.

## Ethics declarations

This study was approved by the Ethics Committee of the Institute of Psychology, Chinese Academy of Sciences (H21077). Informed consent was obtained from all participants involved in the study.

## Consent for publication

Not applicable.

## Notes

### Competing Interest Statement

The authors have declared no competing interest.

